# BeWo-derived extracellular vesicles downregulate IL-6Rα expression via miRNAs on CD4+ T lymphocytes

**DOI:** 10.1101/2025.09.08.674438

**Authors:** Gábor Seregélyes, Bence Nagy, Árpád Ferenc Kovács, Nóra Fekete, Edit I Buzás, Éva Pállinger

**Affiliations:** Institute of Genetics, Cell- and Immunobiology, Semmelweis University, Budapest, Hungary; Heart and Vascular Center, Semmelweis University, Budapest, Hungary; Department of Pathology and Experimental Cancer Research, Semmelweis University, Budapest, Hungary; MTA-SE Immune-Proteogenomics Extracellular Vesicle Research Group, Budapest, Hungary; HCEMM-SE Extracellular Vesicle Research Group

**Keywords:** human pregnancy, immune tolerance, CD4+ T cells, regulatory T cells, extracellular vesicles, interleukin-6, IL-6Rα, IL-6 pathway, miRNA

## Abstract

Regulatory T lymphocytes are essential for maternal immunotolerance. Their de novo differentiation in the placenta is regulated by local intercellular interactions involving primed uterine immune cells, fetal syncytiotrophoblasts, and the cytokine environment. Trophoblast-derived, HLA-G-positive extracellular vesicles (EVs) can bind to T lymphocytes, thereby influencing their differentiation and cytokine production. Thus, these EVs play a role in establishing and maintaining a tolerogenic environment. In our study, we used the BeWo choriocarcinoma cell line to model the effects of trophoblast-derived EVs. Large EVs derived from BeWo cells (BeWo-12.5K lEVs) reduce IL-6Rα expression on CD4+ T cells. This modifies the IL-6 pathway by downregulating the transcription factors STAT3 and NFKB1, and PIAS3, while upregulating STAT1. This may be caused by specific microRNAs (miRNAs), such as hsa-mir-92a-3p, hsa-mir-520f-3p, and hsa-mir-25-3p. These microRNAs are present in BeWo-12.5K lEVs and target the IL-6 pathway. BeWo-12.5K lEVs induce phenotypic and functional changes in T cells, enhancing the ratio of CD4+/CD25+ T cells that produce IL-10. Pregnancy-associated IL-6Rα downregulation has been demonstrated in clinical samples. Significantly lower levels of IL-6Rα were detected on circulating CD4+CD25+ T cells in healthy pregnant women than in healthy non-pregnant individuals. This finding reflects the in vivo significance of our in vitro studies. Our studies suggest that communication between maternal and fetal cells significantly influences the development and maintenance of local T cell polarity. The microRNA content of HLA-G+ 12.5K lEVs appears to be key to this process, as it alters the IL-6 pathway.

## Introduction

The adaptation of the maternal immune system to pregnancy is strictly regulated and can be characterized by immunological stages, or the "immune clock" [1]. Recognizing the semi-allogeneic fetus and developing maternal immune tolerance are prerequisites for a successful pregnancy. Although an inflammatory environment is necessary for embryo implantation, the immune system transitions to an anti-inflammatory state by the end of the first trimester [2]. The presence of regulatory T cells (Treg) and Th2 cells in the placenta is essential for fetal growth [3–6]. Alterations to this immune milieu can result in recurrent spontaneous abortion [7, 8]. The local cytokine environment determines the polarization of naïve CD4+ T helper (Th) cells into specific subsets, such as Th1, Th2, Th17, and Treg cells. Transforming growth factor beta (TGF-β) and interleukin 2 (IL-2) are critical for Treg differentiation. The final effects of TGF-β signaling are also highly influenced by surrounding cytokines. Proinflammatory cytokines, such as IL-1β, IL-6, and TNF-α, can neutralize the effects of TGF-β and favor Th17 development over Treg polarization [9, 10]. IL-6 is a Janus faced player during pregnancy [11]. Although IL-6 is present in the amniotic fluid and the placenta, already from the earliest period of pregnancy [12, 13], the complete role of it has not been clarified yet. According to our knowledge IL-6, as a pleiotropic cytokine contributes to the main physiological processes during pregnancy, including implantation, embryogenesis and parturition [14]. On the other hand, IL-6 directly acts on CD4+ Th cells and plays a role in the orchestration of Th cell differentiation and function [15, 16].

The tolerogenic immune environment at the feto-maternal interface is maintained through direct cell-to-cell interactions, soluble mediators, and extracellular vesicles (EVs) [17, 18]. The presence of human leukocyte antigen-G (HLA-G) -positive trophoblast-derived EVs at the feto-maternal interface and in the maternal bloodstream is a classic feature of pregnancy [19–21]. EVs are membrane-covered intercellular messengers that are constitutively secreted by eukaryotic and prokaryotic cells [22]. Two types of EVs can be distinguished according to their production: ectosomes and exosomes. However, available isolation methods separate them based on size [23]. In this context, the following definitions apply to EV populations: small EVs (sEVs), which have an average diameter under 200 nm and are enriched in exosomes, and large EVs, which have an average diameter higher than 200 nm. The large EV fraction contains the "12.5K lEVs" (formerly known as iEVs or microvesicles), which have been isolated and purified by centrifugation at 12,500 *g*. [24] These EV subpopulations may differ in their membrane components, including proteins and lipids, as well as in their cargo, such as nucleic acids (e.g., microRNAs, intact mRNAs, long non-coding RNAs, and double-stranded DNA) [25]. Placental EVs are critical modulators of the maternal immune system [26]. Because EVs are involved in many physiological and pathological processes, they can serve as diagnostic and prognostic biomarkers for various pathological conditions [27]. In our previous study, we found that both platelet- and trophoblast-derived EVs target immune cells [28] and trophoblastic-derived 12.5K lEVs induced Treg expansion is associated with the HSPE1 expression and the specific miRNA cargo of EVs. [29]

In the present study, we demonstrate that trophoblast-derived EVs modulate the IL-6 sensitivity of Th cells by regulating their interleukin-6 receptor alpha (IL6Ra) expression via miRNA, thereby playing an important role in local Treg cell polarization.

## Material and methods

### Patients and sample collection

Peripheral venous blood samples were collected from the median cubital vein of 37 healthy pregnant women in their 18th week of pregnancy and 15 healthy non-pregnant women using Vacutainer® Brand Plus Sodium Heparin Tubes or Vacutainer® EDTA Tubes from Becton Dickinson (BD, San Jose, California, USA). The blood samples arrived at the laboratory within two hours and were immediately processed. The pregnant women underwent routine obstetrical control at the Department of Obstetrics and Gynecology at Semmelweis University’s Faculty of Medicine. An obstetrical history of previous spontaneous abortions, the course of previous pregnancies, hypertension, gestational diabetes, and preeclampsia were exclusion criteria. Samples from the healthy non-pregnant women were obtained from the Hungarian National Blood Transfusion Service. The clinical data of the studied groups are presented in Supplementary Table 1. The study was approved by the Ethics Committee of Scientific Research of Hungary (ETT-TUKEB: 10147-4/2015/EKU (93/2014)).

### Lymphocyte separation

Peripheral blood mononuclear cells (PBMCs) were isolated using density gradient centrifugation with Histopaque® 1077 (Sigma-Aldrich, St. Louis, Missouri), and the monocytes were removed by adherence.

### Cell culture (BeWo cells and human lymphocyte co-culturing)

BeWo choriocarcinoma cell line (ATCC^®^ CCL-98™) was obtained from the American Type Culture Collection. The BeWo cells were cultured as described earlier [29].

To address the interactions between BeWo cells and lymphocytes we designed a co-culture system: 2 x 10^5^ BeWo cells and 1 x 10^6^ lymphocytes were plated into the wells of a BD Falcon™ 12-well Cell Culture Insert Companion Plates (BD San Jose, California, USA) in 2 mL culture media, for 24 hours. BD Falcon™ Cell Culture Inserts (with 1 µm pores) have been used to study the effects of BeWo-derived EVs and soluble mediators on lymphocytes (Supplementary Figure 1).

BeWo cells were tested regularly for mycoplasma infection with a PCR-based method (Supplementary Figure 2A). We examined the viability of the co-cultured cells regularly using a microscope. A flow cytometric analysis was performed at the end of the culturing period. (Supplementary Figure 2B).

Flow cytometry was used to demonstrate the transport of EVs through 1 µm inserts. (Supplementary Figure 2C).

### Isolation of BeWo-derived large vesicles (12.5K lEVs), small-sized vesicles (sEVs) and EV free supernatant by differential centrifugation

2 x 10^6^ BeWo cells were seeded in 75 cm^2^ tissue culture treated flasks in 10 mL of culture media. Once the cells reached 80-90% confluence, we discarded the culture medium and replaced it with 10 mL of FBS-free medium. Then, we cultured the cells for an additional 24 hours. We applied the differential centrifugation method for EV isolation, which yields intermediate recovery and specificity. Next, BeWo cell-culture supernatant was centrifuged at 300 *g* for 5 minutes at room temperature (RT) to remove cell debris. The obtained supernatant was centrifuged at 2 000 *g* for 15 minutes to remove the apoptotic body fraction. The supernatant of the 2 000 *g* speed centrifugation was sedimented at 12 500 *g* for 20 minutes at 16^◦^C (Z216 MK Microlite centrifuge, fixed angle 200.88 rotor, Hermle Labortechnik GmbH, Wehingen, Germany) and subsequently washed with 0.20 µm filtered phosphate-buffered saline (PBS) at 12 500 *g* for 15 minutes at 16^◦^C. The supernatant of 12.5K centrifugation was further processed at 100 000 *g* for 70 minutes at 4^◦^C (Optima MAX-XP, fixed angle MLA-55 rotor, Beckman Coulter Inc, Brea, USA). A subsequent washing step with filtered (0.20 µm pore size, Millipore) PBS at 100 000 *g* for 70 minutes at 4^◦^C was done. The supernatant of the 100 000 *g* washing step was further centrifuged for another 24 hours to obtain the EV free supernatant fraction. The obtained pellets enriched in 12.5K lEV fraction (12.5K pellet) or sEV fraction (100K pellet) were resuspended in 20-100 µL PBS or serum-free Ham’s F12/K media depending on the downstream analysis (Supplementary Figure 3).

### EV characterization

Following the recommendations of the International Society for Extracellular Vesicles’ (ISEV) latest position paper [24], the isolated EV preparations were characterized by several methods. (Supplementary Figure 4)

### miRNA sequencing

Three biological replicates of BeWo-12.5K lEVs were isolated and resuspended in 12 µL of DNase, RNase free water for miRNA sequencing. The RNA integrity was evaluated with Agilent Tapestation and samples with RIN values ≥7 were further processed. For library preparation multiplex Small RNA Libr. Prep Kit was used according to the manufacturer’s instructions. The sequencing was performed on an Illumina MiSeq instrument and the paired-end read value was > 20 million/sample. Next, we aligned the transcriptome sequence reads to the reference genome (Ensembl GRCh38 release) with STAR version 2.5.3a using 2-pass alignment mode. The mirBase annotation was used in both mapping and read counting. After alignment, the reads were associated with known miRNAs and the number of reads assigned to each miRNA was counted using featureCounts from R package Rsubread. The data were normalised using the TMM normalisation method of the edgeR R/Bioconductor package (R version 3.4.1, Bioconductor version 3.5). After pre-processing and quality control we analysed the target genes of the miRNAs (miRDB target score ≥80) and the IL-6R pathway targeting miRNAs with Funrich 3.1.3 [32], g:Profiler [30] and NetworkAnalyst 3.0 [33]. The miRNA-Seq data has been deposited to the ExoCarta database [34].

### Investigation of target cells of BeWo-12.5K lEVs and characterization of molecular interactions between 12.5K lEVs and lymphocytes

One hundred µL of the separated 12.5K lEVs were labeled with a freshly prepared PKH26 fluorescent dye solution at a final concentration of 5 µM according to the manufacturer’s instructions (Merck, USA). After staining, the labeled vesicles were washed by adding 1 mL of RPMI medium and centrifuging at 12,500 *g* for 15 minutes at 16 °C. We evaluated the effectiveness of staining by flow cytometry. Validation was performed using a mock control (the same staining procedure without adding EV samples) and differential detergent lysis (Supplementary Figure 5). The number of EVs was determined by adding internal counting beads (PKH Reference Beads, Merck, USA). A constant amount of BeWo-derived PKH labelled 12.5K lEVs (1 x 10^6^) were added to immunophenotyped lymphocytes (5 x 10^5^) for 30 min at 4 °C to investigate EV binding. T lymphocyte subsets were defined by CD3+/CD4+ (Th cells), CD3+/CD4+/CD25+ (activated Th and regulatory Th cells) and CD3+/CD8+ (Tc cells) positivity. B cells were characterized by their CD3-/CD19+ expression, and NK cells by their CD3-/CD56+ immunophenotype. Antibodies were titrated for the optimal concentration. At the end of the incubation period, cells were washed with PBS and analyzed by flow cytometry.

To characterize the molecular interactions between EVs and lymphocytes, we measured the binding of PKH-labeled EVs after masking surface PS, CD95L, or HLA-G molecules using annexin V, anti-CD95L, or anti-HLA-G antibodies. On the other hand, the surface phosphatidylserine receptor (PSR) and CD95 molecules on lymphocytes were also blocked with anti-PSR polyclonal antibody or anti-CD95 antibodies. PKH-labeled BeWo-derived 12.5K lEVs and immunophenotyped lymphocytes PKH-labeled BeWo-derived 12.5K lEVs and immunophenotyped lymphocytes were incubated together in the presence or in absence of monoclonal antibodies in PBS containing 4% bovine serum albumin (BSA) for 30 minutes at 4 C.

### Investigation of IL-4, IL-6, IL-17 and IFNγ production by *Cytometric Bead Array (CBA)*

IL-4, IL-6, IL-17 and IFN-γ production was detected from cell culture supernatants, by the BD™ CBA (BD San Jose, California, USA) system according to the manufacturer’s instructions (BD Cytometric Bead Array (CBA) Human Th1/Th2/Th17 Cytokine Kit).

### Intracellular IL-10 staining of CD4+ T lymphocytes

5×10^5^ human lymphocytes were stimulated with BeWo-12.5K lEVs for 24 hours in the presence of the protein transport inhibitor Brefeldin A (Thermo Fisher Scientific, USA). Stimulated cells were washed and incubated in 100 µL of staining buffer (PBS containing 1% BSA) with a pre-titrated optimal concentration of PerCP-conjugated anti-human CD4 monoclonal antibody (15 minutes at RT on ice). The immunophenotyped cells were fixed in a 4% paraformaldehyde solution (Sigma-Aldrich Co., St. Louis, MO) for 10 minutes at RT, permeabilized with a 0.1% saponin solution, and incubated in 50 µL of staining buffer (1% FBS containing PBS) containing a pre-titrated optimal concentration of phycoerythrin-conjugated anti-human IL-10 monoclonal antibody for 20 minutes at RT on ice. The cells were washed in 2 mL of PBS and centrifuged at 300 *g* for five minutes. The pellet was resuspended in 300 µL of 2% PFA and analyzed by flow cytometry.

### Study of IL-6R**α** expression on lymphocytes and BeWo cells

We examined the expression of IL-6Rα in lymphocytes obtained from co-culture systems through immunophenotyping. Staining was performed according to the manufacturer’s instructions. A total of 5,000 cells were analyzed per tube.

Adherent BeWo cells were scraped off by cell scraper after a 20 minutes incubation period on ice and were stained using the same staining method described for lymphocyte immunophenotyping.

### Detection of soluble IL-6R**α**

Human Instant ELISA™ (Thermo Fisher, USA) was used for detection of soluble IL-6Rα concentrations according to the manufacturer’s instructions. IL-6Rα levels were detected in pg/mL at 450 nm and 620 wavelengths by Multiskan MS Microplate Reader (Thermo Fisher, USA). The sensitivity of the kit was 0.01 ng/mL.

### IL-6 pathway signalling

For gene expression analysis, first total RNA isolated from untreated and BeWo-12.5K lEVs-treated lymphocytes was purified with RNeasy Mini Kit (Qiagen, CA, USA). The RNA concentration of each sample was determined using the Qubit RNA HR Assay Kit (Qubit Fluorometer, Life Technologies). Samples were stored at −80°C until cDNA synthesis. From each sample 600 ng total RNA was reverse transcribed with Sensifast cDNA Synthesis Kit (Bioline, London, UK). Quantitative PCR reactions using SensiFAST SYBR Hi-ROX Kit with SYBR Green primers (Supplementary table 2) were carried out on an ABI 7900 real-time PCR instrument (Thermo Fisher Scientific, USA) according to the manufacturer’s instructions. Real-time PCR results were calculated according to the following protocol: relative expression level = 2 ^-ΔCt^, where ΔCt = Ct (of gene of interest) – Ct (of houseskeeping gene) with *HPRT* as an endogenous control.

To establish the optimal timepoint for the gene expression analysis we quantified the mRNA levels of key IL-6 pathway factors (*IL6, IL6R, IL6ST, AKT1, STAT1, STAT3* and *PIAS3*) in lymphocytes at 2, 4, 6, 8 and 24 hours after recombinant IL-6 (Sigma-Aldrich Co., St. Louis, USA) (0.02 ng/ mL = 5.0 × 10^7 units/ mg) treatment. The concentration of IL-6 was chosen accordingly to physiological circulating IL-6 concentration in healthy pregnancy [35, 36]. Based on the results of this experiment, we chose the four-hour treatment as the optional time point for IL-6 pathway analysis.

### Flow cytometry

Antibodies and dies used in flow cytometry are listed in Supplementary Table 3. Measurements were carried out using a FACSCalibur flow cytometer (BD, San Jose, CA, USA) on the day of the staining. In the case of cellular measurements, 5×10^4^ cells/ tube were collected. Data were analysed with CellQuestPro (BD, San Jose, California, USA) and Flowjo v10.3 (Tree Star Inc., OR, USA) software.

### Statistics

GraphPad Prism version 8.0 (GraphPad Software, La Jolla California, USA) was used for statistical analysis. For normally distributed data Two-sided Student’s unpaired t-test, respectively for multiple parameter analysis ANOVA test was used, followed by Bonferroni correction. For data not showing normal distribution Mann-Whitney U or Wilcoxon test was used, for multiple parameter analysis ANOVA test was used, followed by Kruskal-Wallis test. The level of significance was set at p <0.05.

The datasets generated during and/or analysed during the current study are available from the corresponding author on reasonable request.

## Results

### Target cells of BeWo-EVs

The 12.5k lEVs primarily bound to CD3+ T lymphocytes; however, the binding ability of T cell subsets differed. BeWo-lEVs preferred Th cells (5.04 ± 0.98 %) compared to Tc lymphocytes (2.01 ± 0.14 %) (p < 0.05). On the other hand, significantly fewer CD19+ B cells (0.92 ± 0.45 %, p < 0.01) and CD56+ NK cells bound BeWo-lEVs (0.14 ± 0.05 %) (p < 0.001) like T lymphocytes (Figure 1A-B). There was no difference in the binding ability to the lymphocytes of pregnant and non-pregnant donors (non-pregnants: 3.55 ± 0.48 % vs. pregnant: 5.04 ± 0.98 %) (p>0.05). We used monoclonal antibodies or AnnexinV pretreatments for blocking the BeWo-lEVs - lymphocyte interactions (Figure 1C-D). Anti-phosphatidylserine receptor (PSR) mAb pre-treatment of lymphocytes partially inhibited the binding of 12.5K lEVs to CD3+/CD4+ (untreated vs. anti-PSR Mab: 215 ± 106 mean fluorescence intensity (MFI) vs. 112.5 ± 24.06 MFI) and to CD3+/CD8+ T cells (untreated vs. anti-PSR Mab: 33.5 ± 9.3 MFI vs. 21.43 ± 7.17 MFI). Similarly, Annexin V pre-treatment of BeWo-lEVs resulted in minimal binding inhibition (untreated vs. Annexin V treated: 13.2 ± 0.46 MFI vs. 7.06 ± 0.4 MFI). Masking of HLA-G on BeWo-lEVs (untreated vs. anti-HLA-G mAb: 13.05 ± 0.55 MFI vs. 7.83 ± 0.06 MFI) could also prevent the binding of 12.5K lEVs to lymphocytes. Neutralization of CD95 molecules on lymphocytes (untreated vs. anti-CD95L mAb: 43.53 ±1.88 MFI vs. 39.5 ± 4.8 MFI) or CD95L (untreated vs. anti-CD95 mAb: 13.05 ± 0.55 MFI vs. 16.35 ± 0.35 MFI) on 12.5K lEVs did not have a significant effect on the 12.5K lEV-lymphocyte interaction. Based on these results we concluded, that interactions via PS and HLA-G could coordinate, at least partially the binding of 12.5K lEVs to CD4+ lymphocytes (Figure 1C-D).

**Figure 1.**
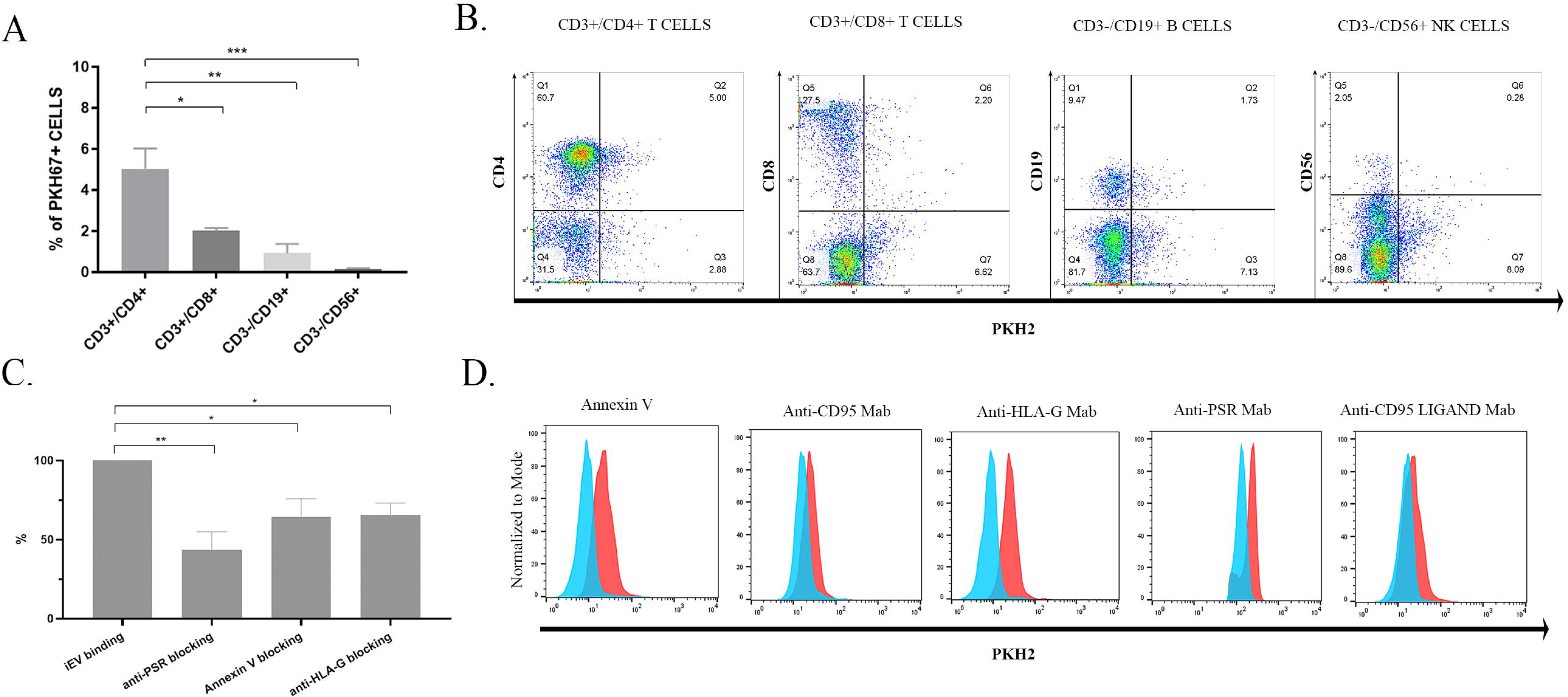
Target cells of BeWo-derived 12.5K lEVs A. Binding of BeWo-12.5K lEVs to immunophenotyped lymphocytes isolated from pregnant women BeWo-12.5K lEVs bound significantly more CD3+/CD4+ T cells (n= 6 pregnant donors, one-way ANOVA followed by Bonferroni correction). B. Representative dot plots show the cell-bound 12.5K lEVs to the different lymphocyte subpopulations. C. Analysis of selected blocking between 12.5K lEVs and their target CD4+ T cells. Anti-PSR, annexin V and anti-HLA-G blocking resulted in significant inhibition (n=6 pregnant donors, one-way ANOVA followed by Bonferroni correction). D. Representative histograms show the changes of expression levels after selective blocking of a specific interaction between 12.5K lEVs and target lymphocytes (red histogram: PKH2+ surface bound 12.5K lEVs, blue histogram: PKH+ surface bound 12.5K lEVs after targeted blocking, n= 6 different pregnant donors; measurements were repeated 2 times from each donor).

### The effects of BeWo-lymphocyte interactions on the IL-6R**α** expression of T cells

IL-6Rα expressing cells were detected in all of the tested T cell subsets, though the proportion of IL-6Rα+ cells varied among them (CD4+ T cells: 61.8 ± 6.3%; CD8+ T cells: 11.4 ± 1.9%). Since IL6 sensitivity depends on the number of IL-6Rα, we compared their expression levels in T cell subsets. Exofacial IL-6Rα was significantly higher on CD4+ T cells (13.72 ± 0.68 MFI) than on CD8+ T cells (9.31 ± 0.16 MFI). (p < 0.0001) (Supplementary Figure 6A-D).

We examined the impact of BeWo-12.5K lEVs on the IL-6Rα expression of T lymphocytes in BeWo-lymphocyte co-culture systems. BeWo-derived 12.5K lEVs significantly downregulated the exofacial IL-6Rα on lymphocytes (Ly + 12.5K lEV: 9.7 ± 0.06 MFI) (p < 0.05) (Figure 2A-B). Since T lymphocytes were identified as targets of BeWo-12.5K lEVs, the IL-6Rα levels of CD4+ T cell subsets were further analysed. The Th cells (untreated: 13.71 ± 0.68 MFI; co-cultured: 9.2 ± 0.04 MFI) and the CD25+ Th subset (untreated: 16.52 ± 1.18 MFI; co-cultured: 9.49 ± 0.27 MFI) exhibited significantly lower IL6Ra expression levels when the co-cultured cells were separated by insert membranes (p<0.0001) (Supplementary Figure 6 E-F). Direct cellular interaction (Ly + BeWo) did not have any effect (data not shown). Soluble IL-6Rα can be formed by ectodomain shedding or alternative splicing, but cells can also lose their exofacial IL-6Rα via vesicle-associated shedding [37]. To determine the pathways of IL-6Rα shedding, the concentration of soluble IL-6Rα and also the EV-bound IL-6Rα was determined by ELISA or flow cytometry, respectively. Treatment of lymphocytes with BeWo-derived 12.5K lEVs did not change either the soluble IL-6Rα concentration in cell culture supernatant (untreated vs. 12.5K lEV treated: 114 ± 45.8 pg/ mL vs. 178.3 ± 24.1 pg/ mL p>0.05) or the number of IL-6Rα positive 12.5K lEVs (untreated vs. 12.5K lEV treated: 741 ± 326 events/ µL vs 1071 ± 351 events/ µL, data not shown), which may suggest that the IL-6Rα downregulation could happen at the gene expression level.

**Figure 2.**
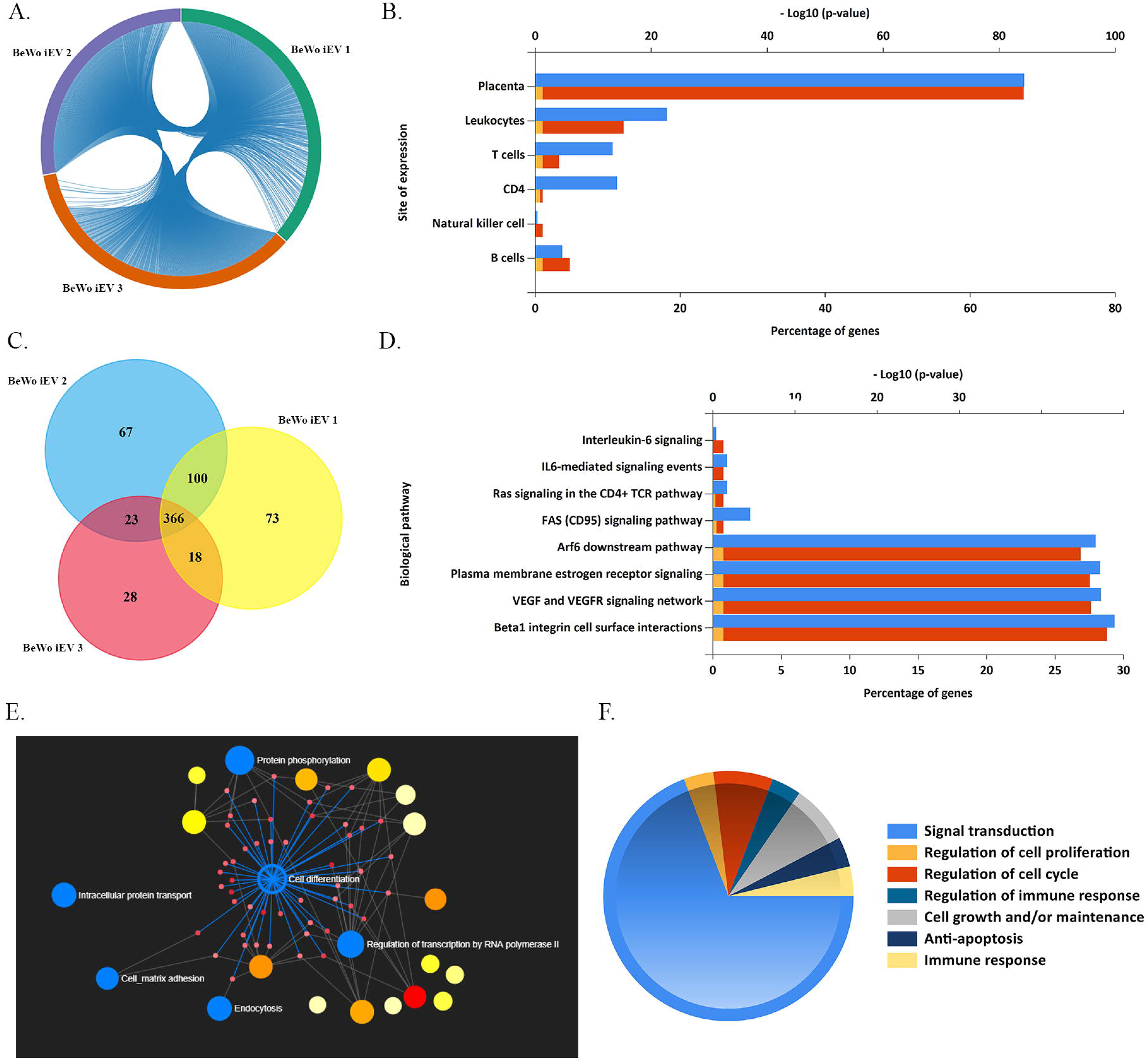
IL-6Rα downregulation mediated by miRNAs A. Analysis of IL-6 downregulation upon BeWo-12.5K lEV treatment (n=10 donors). B. Representative FACS histogram showing the IL-6Rα downregulation upon 12.5K lEV treatment marked with red (untreated control - blue) C. Molecular function analysis of BeWo-derived miRNAs target genes revealing the interleukin-6 receptor binding as one of the main targets. D. Overall, we identified 366 common miRNA in 12.5K lEVs. 21 miRNAs are targeting the downregulation of IL-6R, together with the expression values (RPM). E. IL-6R downregulating miRNA network with highlighted (dark blue) miRNA found in the BeWo-12.5K lEVs. Values represent mean ± SEM (n=10 ** p < 0.01, *** p < 0.001) Ly - lymphocytes

To investigate the mechanism of IL-6 downregulation we performed miRNA-Seq of BeWo-derived 12.5K lEVs. Altogether we identified 517 ± 70 miRNAs in the 12.5K lEV samples, out of which 366 were common in all samples. The biological pathway analysis revealed that the interleukin-6 receptor binding (GO:0005138) to be significantly enriched (p_adj_ = 0.02) (Figure 2C) and 21 miRNAs from the 366 were targeting IL-6R. Overall, in the NetworkAnalyst database there are 103 known miRNAs targeting IL-6R, through gene silencing of *IL6ST* and *ERAP*. The top 5 highly expressed miRNAs targeting IL-6R found in BeWo-derived 12.5K lEVs were the hsa-miR-92a-3p (18170 ± 5221 RPM), hsa-miR-520f-3p (5274 ± 1002 RPM), hsa-miR-25-3p (5065 ± 838.8 RPM), hsa-miR-519d-3p (3703 ±1261 RPM) and hsa-miR-26b-5p (1348 ± 363.3 RPM) (Figure 2D-E).

Further analysis of the miRNA pattern revealed that there is a strong miRNA signature pattern among the biological replicates (Figure 3A and C). Most of the target genes of the miRNAs found in BeWo-12.5K lEVs are expressed in the placental tissue and one of the main target cell types is the CD4+ T cell (Figure 3B). Beside the IL-6 signalling pathway, other significantly enriched targeted biological pathways were FAS and Arf6 downstream pathways, the plasma membrane estrogen receptor and VEGF signalling, followed by the β1integrin cell surface interaction biological pathway (Figure 3D). Additionally, BeWo-12.5K lEVs induced molecular function network analysis reveals cell differentiation as one of the main significant molecular function (p=0.001, Figure 3E). The in-silico analysis of cell differentiation depicts that the process is initiated by signal transduction (IL6R downregulation) followed by regulation of cell proliferation and cell cycle coupled with anti-apoptotic signalling events. Furthermore, cell growth and immune response balancing accompany cell differentiation (Figure 3F).

**Figure 3.**
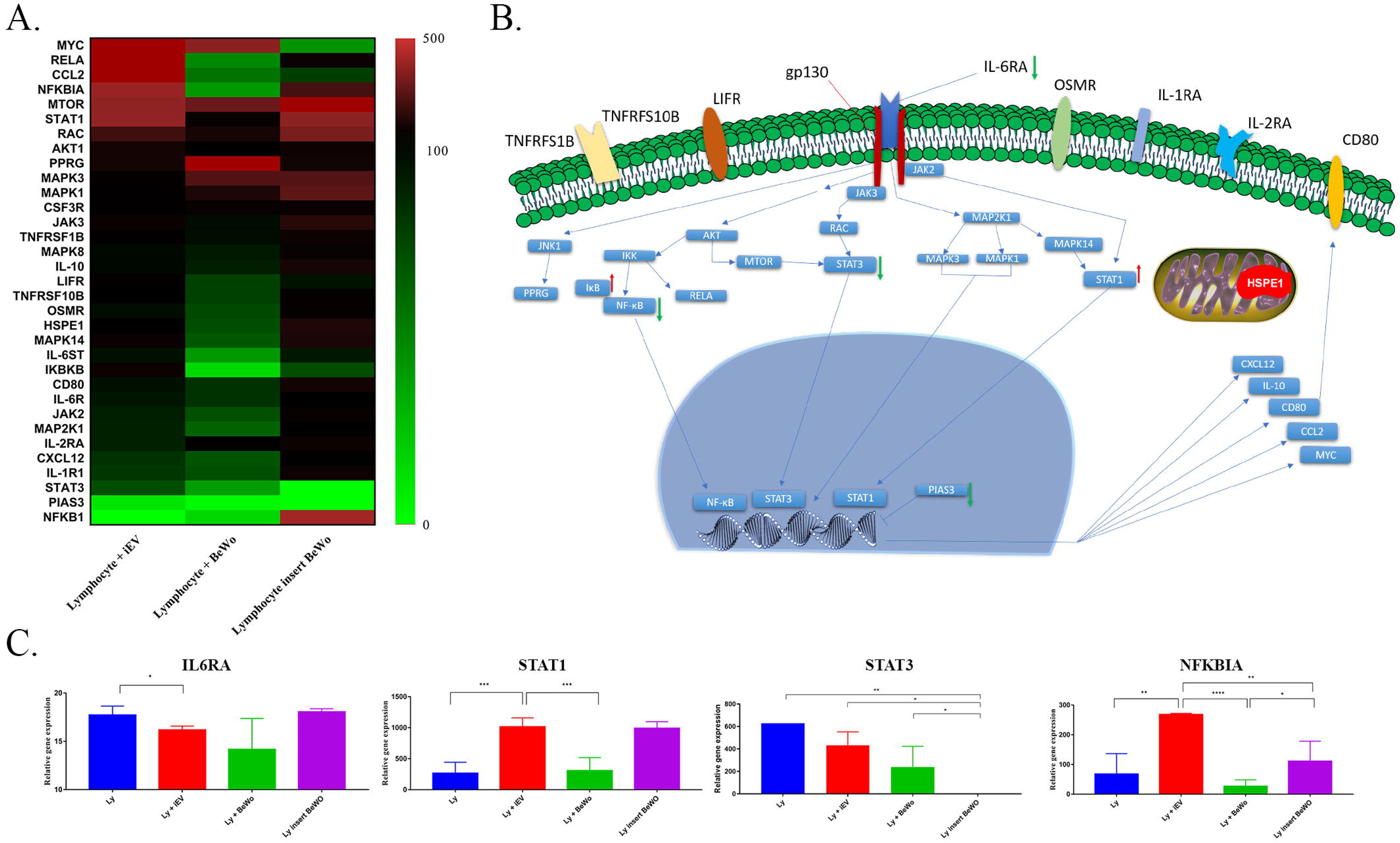
BeWo-derived 12.5K lEV miRNA content A. Chord diagram analysis showing the overlap between the biological replicates. B. Bar chart showing the site expression of genes targeted by BeWo-12.5K lEVs. Based on the analysis the main target tissue is the placenta. One of the prominent cell types are the leukocytes, most of the targeted cells are the CD4+ T cells (blue bars represent the percentage of target genes, red bars are representing the p-values, and the yellow bars represent the p=0.05 reference). C. Venn diagram showing the 366-common miRNA in 12.5K lEVs. D. Biological pathway analysis showing the miRNA targeted pathways, among the IL-6 signalling, the following pathways are significantly targeted: FAS signalling and Arf6 downstream pathways, the plasma membrane estrogen receptor and VEGF signalling. E. BeWo-derived 12.5K lEV induced molecular function network analysis reveals as the main molecular function the cell differentiation. F. The in-silico detailed analysis of cell differentiation function reveals the molecular function is mainly induced through alteration of signal transduction (IL-6R downregulation), followed by regulation of cell proliferation and cell cycle coupled with an anti-apoptotic signalling event. Furthermore, cell growth and immune response balancing accompany the cell differentiation.

### IL-6 pathway analysis

Lymphocytes pre-treated with 12.5K lEVs and further treated with recombinant IL-6 showed a decreased *IL6R* expression. *STAT1* showed a 3.5-fold change increase in mRNA expression upon 12.5K lEVs treatment (relative gene expression of untreated: 265.8 ± 125.8, 12.5K lEVs treated: 1014 ± 102.4, Ly + BeWo: 308.4 ± 148.3, Ly insert BeWo: 990.8 ± 75.2). *PIAS3* showed a 10 fold decrease in 12.5K lEVs treated lymphocytes (relative gene expression of untreated: 44.7 ± 0.07, 12.5K lEVs treated: 4.48 ± 0.017, Ly + BeWo: 1.09 ± 0.37, Ly insert Bewo: not detected), *AKT1* showed an 1.4 fold increase upon 12.5K lEVs stimulation (relative gene expression of untreated: 16.8 ± 0.68, 12.5K lEV treated: 23.3 ± 0.4, Ly + BeWo: 17.39 ± 3.15, Ly insert BeWo: 21.15 ± 4.0), *RELA*, *NFKBIA* showed an increased mRNA expression (Figure 4). In BeWo-lymphocyte co-culture system we detected a decrease in *CXCL12*, *IL1R1*, *IL6ST*, *OSMR*, *TNFRSF10B* and *STAT3* expression. Separation of BeWo cells and lymphocytes with 1 µm pore sized membrane led to increased *IL10*, *IL2RA*, *IL1R1*, *CD80*, *JAK2*, *JAK3*, *MAPK1*, *MAPK14*, *RAC1* and decreased *STAT3* expression detectable in lymphocytes (Supplementary Table 4).

**Figure 4.**
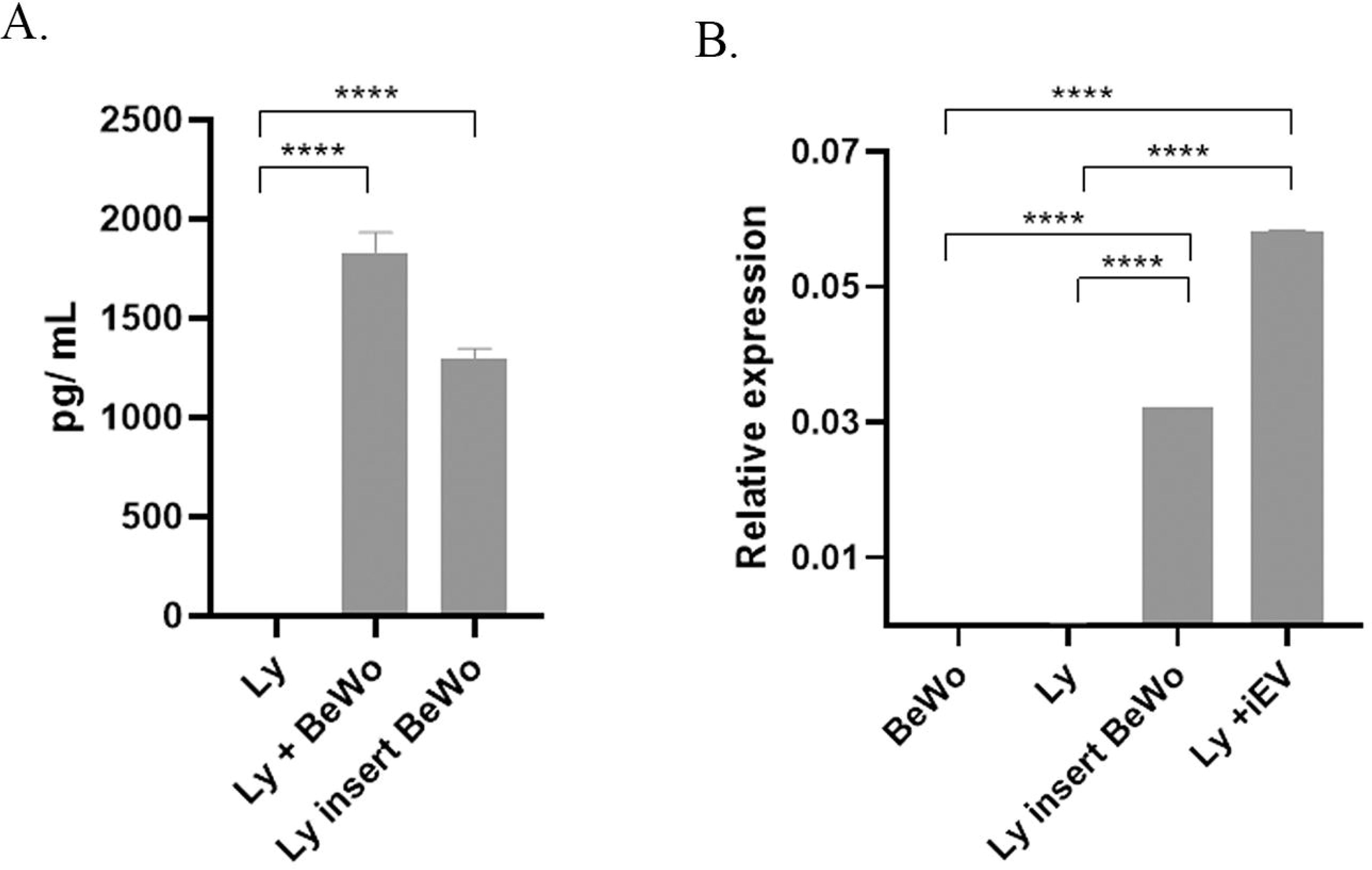
IL-6 pathway analysis A. Heatmap analysis of IL-6 pathway of non-pregnants’ lymphocytes treated with BeWo-12.5K lEVs, lymphocytes co-cultured with BeWo cells with or without a 1µm pore sized insert membrane (n=3, each sample was run in triplicates). Heatmap showing the differential expressed genes of 12.5K lEV treated or BeWo co-cultured lymphocytes normalized to the untreated control lymphocytes. B. Schematic figure of the analysed IL-6 pathway molecules. Green arrows show significant downregulation at mRNA level upon BeWo-derived 12.5K lEV treatment. Red arrows mark the upregulated molecules upon 12.5K lEV treatment. C. Figures showing the relative gene expression: down- or upregulation of *IL6R*, *STAT1, STAT3* and *NFKBIA* mRNAs upon different treatments of lymphocytes (one-way ANOVA followed by Bonferroni correction, * p < 0.05, ** p < 0.01, *** p < 0.001, ****p < 0.0001).

### IL-6 production induced by BeWo-lymphocyte interaction or BeWo-derived 12.5K lEVs

Neither the BeWo cells nor the lymphocytes secreted IL-6 without stimulation; however, both populations began producing it after co-culturing. Higher levels of IL-6 could be measured after direct cell-to-cell interaction than with the transwell system (direct contact: 1,833 ± 30.02 pg/mL and 1,296 ± 15.61 pg/mL, p < 0.001). IL-6 production in the co-culture system was also validated at the mRNA level by qPCR. (Figure 5.).

**Figure 5.**
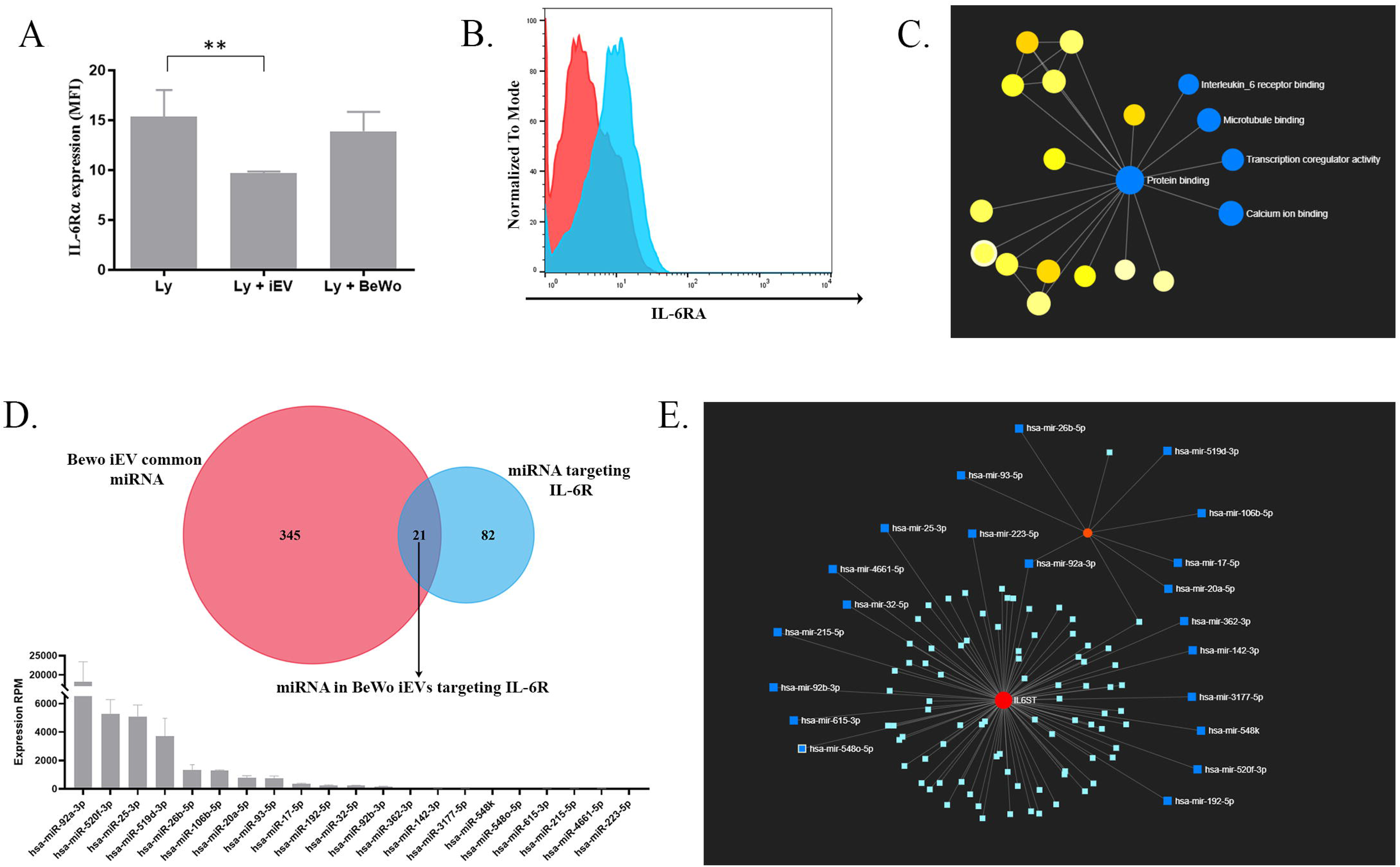
Interleukin 6 secretion A. IL-6 could be detected both in the cell culture supernatant of insert separated BeWo co-culture system and also in the media of BeWo-12.5K lEV treated non-pregnants’ lymphocytes. Unstimulated lymphocytes did not secrete IL-6 (n=11, **** p < 0.0001, One-Way ANOVA, followed by Bonferroni’s correction). B. Validation of IL-6 secretion at the mRNA level.

### IFN-γ, IL-4, IL-10 and IL-17 production induced by BeWo-lymphocyte interactions

Neither the direct cellular interaction nor the membrane separated co-culture system of lymphocytes and BeWo cells induced detectable levels of IFN-γ and IL-4. In the case of IL-17, there was no significant difference compared to control lymphocyte secretion levels (Ly: 58 ± 0.01, Ly insert BeWo: 61.44 ± 0.27; Ly + BeWo: 61.77 ± 0.02, data not shown). However, BeWo-derived 12.5K lEVs induced a significant IL-10 production in CD4+ T cells. To elicit the specific potential of 12.5K lEVs induced IL-10 production, lymphocytes were treated with BeWo-derived 12.5K lEVs, sEVs or the EV-free supernatant. Only the 12.5K lEV induced increased IL-10 expression in CD4+ T cells (12.5K lEVs treatment 305.9 ± 29.1 MFI; sEV treatment: 252 ± 9 MFI; EV-free supernatant: 258.5 ± 45.5 MFI; non-treated CD4+ T cells: 255.3 ± 46.2 MFI, p < 0.01 Wilcoxon test), as determined by multicolour flow cytometry. (Supplementary figure 7.).

### IL-6R**α** expression of circulating lymphocytes of pregnant and non-pregnant women

As our in vitro studies showed a strong IL-6Rα downregulation in CD4+ lymphocytes, we aimed to assess the *in vivo* significance of IL-6Rα downregulation during pregnancy. Peripheral blood samples from first trimester healthy pregnant and non-pregnant control women were assessed for IL-6Rα expression on circulating lymphocytes. We detected the significantly lower percentage of IL-6Rα expressing CD3+/CD4+/CD25+ lymphocytes in the blood of pregnant women compared to the healthy non-pregnant group (81.53 ± 4.2 % vs. 92.78 ± 1.95 %; p < 0.05) (Figure 6 A-D).

**Figure 6.**
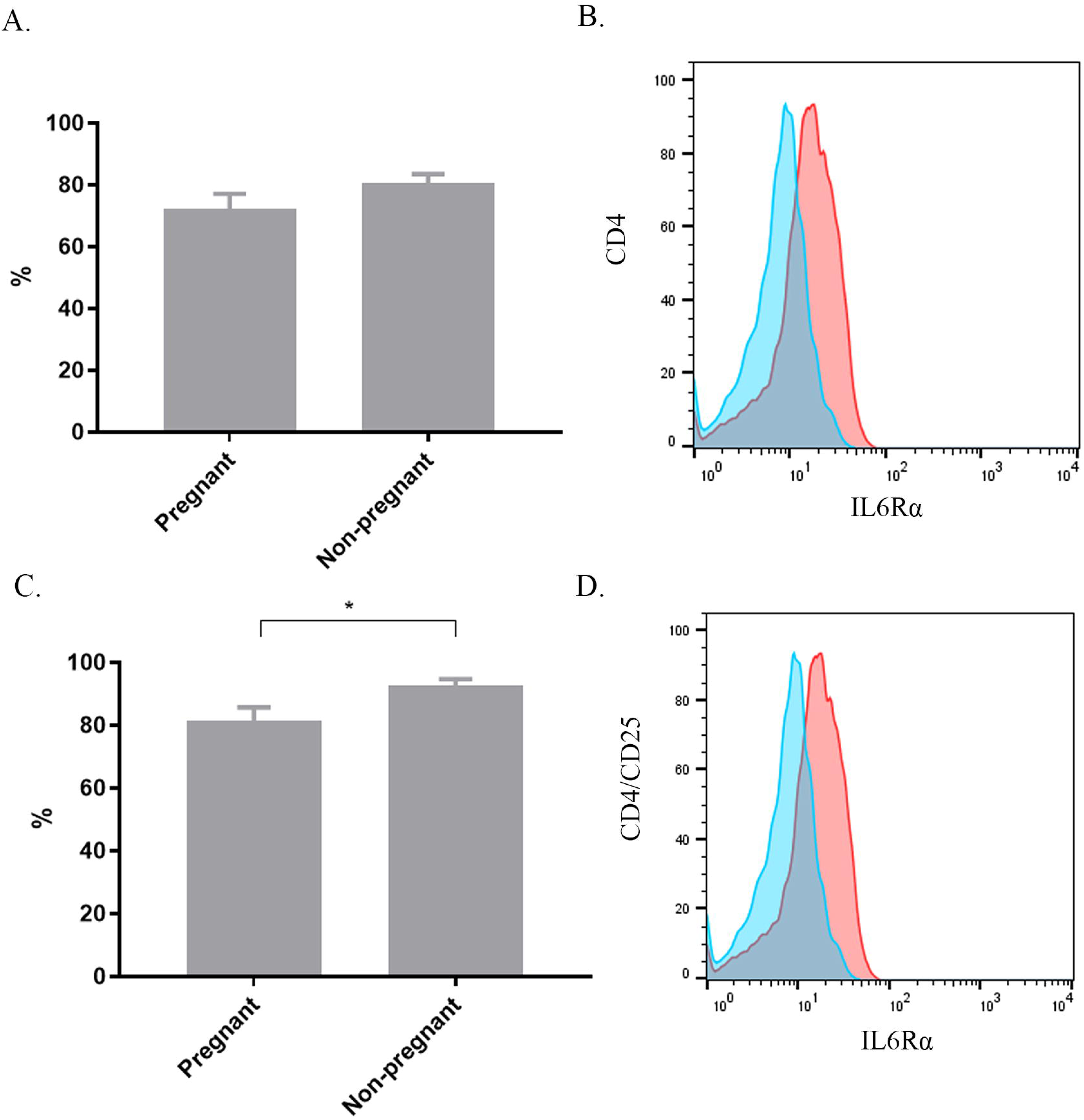
IL6-Rα expression on pregnant and non-pregnant lymphocytes A. Analysis of IL-6Rα expressing circulating CD4+ T cells in healthy non-pregnant and pregnant donors (n=12). B. Representative overlay histograms show the IL-6Rα expression on CD4+ lymphocytes of a pregnant (blue histogram) and a non-pregnant donor (red histogram). C. Percentage of IL-6Rα expressing CD4+/CD25+ T_reg_ cells in healthy non-pregnant and pregnant donors (n=12). D. Representative overlay histograms show the IL-6Rα expression on CD4+/CD25+ lymphocytes of a pregnant (blue histogram) and a non-pregnant donor (red histogram) (unpaired t-test, * p < 0.05).

## Discussion

Local immune tolerance is essential for a successful pregnancy. This antigen-specific immune suppression is mediated by cellular and humoral factors, as well as the recently recognized 12.5K lEV-related network. Through their complex molecular cargo, EVs may play a role in maintaining local immune tolerance. The existence of suppressor regulatory T lymphocytes was revealed in the early 1980s. CD4+/CD25+/FoxP3+ natural T_reg_ cells are characterized by their low proliferating ability, anergy and the production of TGF-β and IL-10 [38, 39]. Although the exact process of T_reg_ cells development is still unknown, peripheral differentiation seems to be an existing pathway beside thymic differentiation [40].

The presence of T_reg_ cells can be shown in the feto-maternal interface during the entire pregnancy and increased percentage of circulating CD4+/CD25+ Treg cells has been found in the blood of healthy pregnant women; however, their level is dependent on gestational age: the highest T_reg_ number can be measured during the second trimester; and their number significantly decreases in 6-8 weeks after delivery [41]. An elevated level of T_reg_ cells can also be detected in local lymph nodes and in vaginal secretions even in the earliest gestational weeks. The question arises whether this early presence of regulatory T cells in pregnant tissues would be a consequence of local differentiation from naïve T cells. One of the theories suggests that fetal antigens presented by local professional antigen presenting cells induce the differentiation of CD4+/CD25- Th lymphocytes into CD4+/CD25+ T_reg_ cells [42, 43].

It is well known that trophoblast cells present fetal alloantigens, playing a role in the induction and maintenance of local immune tolerance at the feto-maternal interface. The role of HLA-G in inducing Treg cells has also been described. In our earlier studies, we demonstrated that HLA-G-expressing, trophoblast-derived 12.5K lEVs, which were isolated from maternal plasma, bind to T lymphocytes. This binding inhibits the IL-2-induced signal transduction pathway of T cells [28]. In the present work, we examined the role of trophoblastic cells and their 12.5K lEVs in the putative local development of Treg cells by using the BeWo choriocarcinoma in vitro model system. Co-culturing of BeWo cells and lymphocytes resulted in IL-6 production. In the present work, we investigated the role of trophoblastic cells and their 12.5K lEVs in the putative local development of Treg cells by using the BeWo choriocarcinoma in vitro model system. Co-culturing of BeWo cells and lymphocytes resulted in IL-6 production. IL-6 has been shown to be a critical cytokine that controls the transition between the induction of regulatory T lymphocytes and Th17 cells [44]. This shift is inauspicious for the pregnancy outcome, so it is cardinal that the immune system counterbalances this effect. The modulation of cytokine-induced cellular effects via regulation of receptor expression is a highly sensitive control mechanism. We determined the IL-6Rα expression of the lymphocytes in order to check their IL-6 sensitivity. BeWo-derived 12.5K lEVs downregulated IL-6Rα expression of CD4+ T lymphocytes, indicating the decreased IL-6 sensitivity. Decrease of IL-6Ra resulted in profound functional changes of lymphocytes including disrupted IL-6 signalization and altered cytokine production. Regulation of cell surface receptor expression can occur through multiple pathways including ectodomain shedding [45], vesicle-mediated removal [46] or transcriptional and posttranscriptional modification [47]. Ectodomain shedding or vesicle-mediated receptor removal could be excluded based on our experiments, therefore, we focused on transcriptional regulation. We premised that the regulation happens by the alteration of the IL-6 pathway, and consequently, the mRNA level of *IL6R* would be downregulated, which we have confirmed by qPCR analysis. Additionally, we observed the downregulation of key players of the IL-6 pathway, including the central component *STAT3* transcription factor. To elicit the downregulation mechanism of *IL6R*, we analysed whether miRNAs are mediating the observed process. miRNAs represent a class of small, noncoding RNA molecules which have major role in the posttranscriptional regulation of protein expression [48]. We employed next-generation sequencing to analyse miRNA profiles in BeWo-12.5K lEVs and identified 21 miRNAs targeting downregulation of *IL6R* through gene silencing.

Our findings indicate that HLA-G expressing trophoblastic 12.5K lEVs have a central role in the regulation of local Th cell polarization via regulation of their IL-6 sensitivity. The appearance of placental EVs in maternal circulation raises the possibility that they play a role in the maintenance of maternal immune tolerance at the systemic level as well. The lower circulating level of IL-6Rα bearing CD4+/CD25+ T cell subset in healthy pregnant women also supports this hypothesis.

It should be noted that in the placental microenvironment, EV-mediated interactions between trophoblast cells and lymphocytes orchestrate the local immune response, which is critical for the maintenance of maternal immune tolerance during pregnancy.

## Supporting information

https://docs.google.com/document/d/1KXazgThKnEBpWNiw_TCeR6llI-KYDagH/edit?usp=drive_link&ouid=102400073143219040591&rtpof=true&sd=true

## Acknowledgement

This work was funded by STIA_2024 (19279/PMKP/2025) to ÉP, Hungarian National Research, Development and Innovation Office—NKFIH, OTKA PD_21 138521 grant to ÁFK. We are grateful for the electronmicroscopic measurements of Tamás Visnovitz.

## Author contribution

BN, GS, NF and ÁFK carried out the experiments; ÁFK, BN, GS and ÉP participated in data analysis; ÁFK, EIB, and ÉP wrote the manuscript; ÉP designed the experiments.

## Abbreviations

EV: extracellular vesicles
12.5K lEV: intermediate-sized extracellular vesicles
FSC: Forward Scatter
HLA-G: human leukocyte antigen G (major histocompatibility complex, class I, G)
IL-6Rα: interleukin 6 receptor alpha
miRNA: micro ribonucleic acid
PS: phosphatidylserine
PSR: phosphatidylserine receptor
sEV: small-sized extracellular vesicles
SSC: Side Scatter

