## Supplementary material for "BeWo-derived extracellular vesicles downregulate IL-6Rα expression via miRNAs on CD4+ T lymphocytes": https://docs.google.com/document/d/1KXazgThKnEBpWNiw_TCeR6llI-KYDagH/edit?usp=drive_link&ouid=102400073143219040591&rtpof=true&sd=true

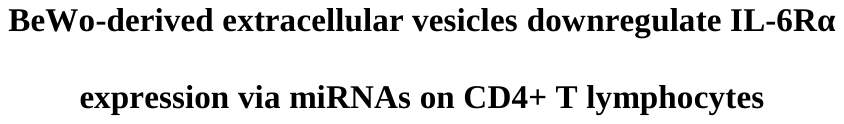

Gábor Seregélyes^1^, Bence Nagy^1^, Árpád Ferenc Kovács^2,3^, Nóra Fekete^1^, Edit Irén Buzás^1,4,5^, Éva Pállinger^1^

^1^ Institute of Genetics, Cell- and Immunobiology, Semmelweis University, Budapest, Hungary

^2^ 1Heart and Vascular Center, Semmelweis University, Budapest, Hungary

^3^Department of Pathology and Experimental Cancer Research, Semmelweis University, Budapest, Hungary

^4^MTA-SE Immune-Proteogenomics Extracellular Vesicle Research Group, Budapest, Hungary

^5^ HCEMM-SE Extracellular Vesicle Research Group

### Supplementary Figures and Tables

#### 1.1 Supplementary Figures

Supplementary Figure 1.

*In vitro* co-culture system to study the interaction between BeWo cells and lymphocytes.

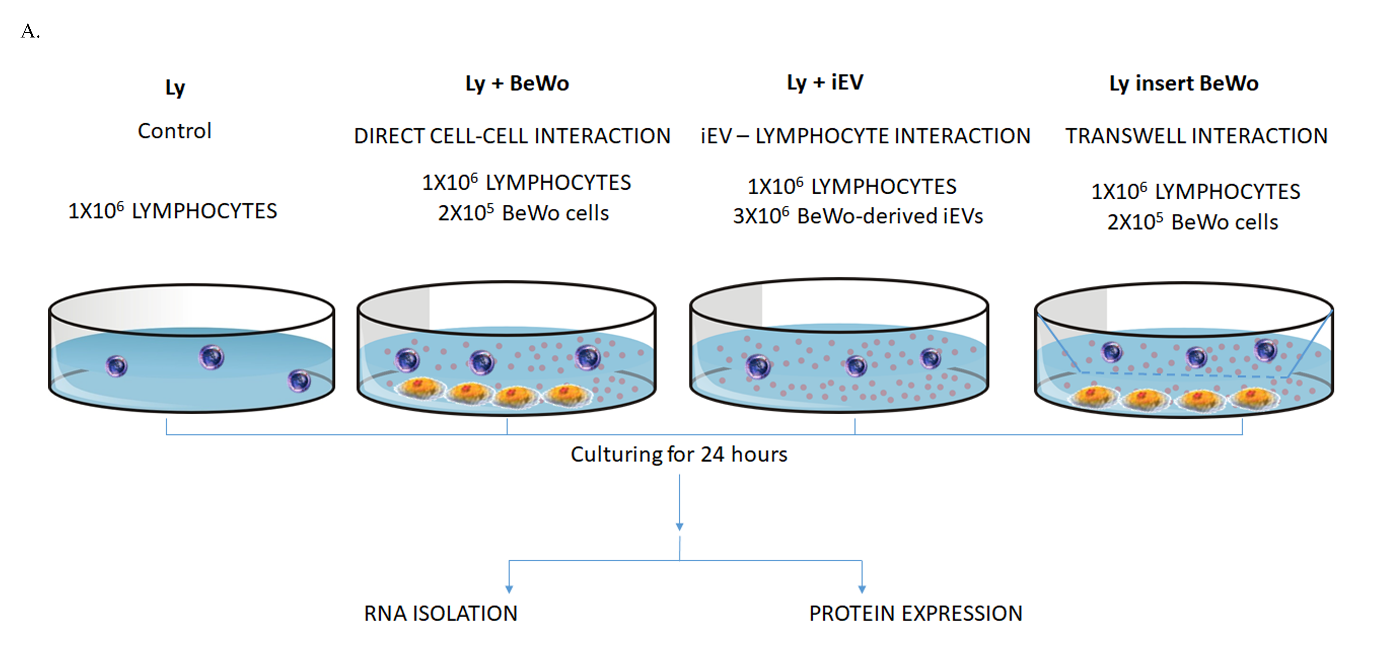

2 x 10^5^ BeWo cells and 1 x 10^6^ lymphocytes were plated in each well of a BD Falcon™ 12-well cell culture insert companion plate (BD, San Jose, CA, USA). The cells were cultured in 2 mL of conditioned culture medium for 24 hours. BD Falcon™ cell culture inserts with 1 µm pores were used to study the effects of BeWo-derived EVs and soluble mediators on lymphocytes.

**Supplementary Figure 2.**

Studies of BeWo cells and insert permeability

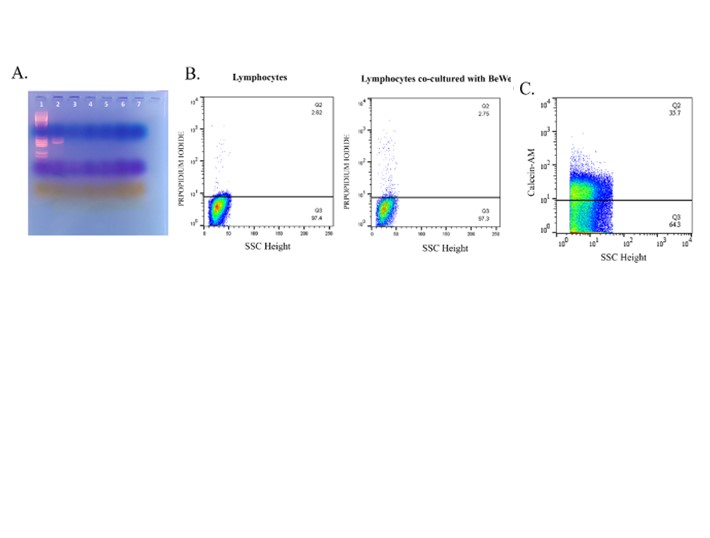

A) BeWo cells were tested regularly for mycoplasma infection by PCR. The results of the PCR-based Mycoplasma test on the BeWo cells are shown in the image below. The image contains a DNA ladder (1), a positive control (2), and the results of the BeWo test (3–7). The test results indicate that there is no mycoplasma contamination.

B. The viability of the BeWo cells was analyzed using flow cytometry to detect propidium iodide (PI) incorporation. Representative dot plots show the PI fluorescence of the cells. In both conditions, more than 95% of the cells were viable (propidium iodide negative).

C. To study the membrane permeabilization of BeWo-derived EVs, we loaded BeWo cells with a calcein stain. The Calcein-labeled BeWo cells were cultured in a cell culture insert plate with a 1 µm pore size. The cell culture supernatant was collected from the upper compartment, and the calcein-stained EVs were detected by flow cytometry. The presence of calcein-stained 12.5K lEVs confirmed that the 12.5K lEVs could freely transfer through the membrane.

**Supplementary Figure 3.**

Isolation of EVs from the serum-starved BeWo cell culture supernatant.
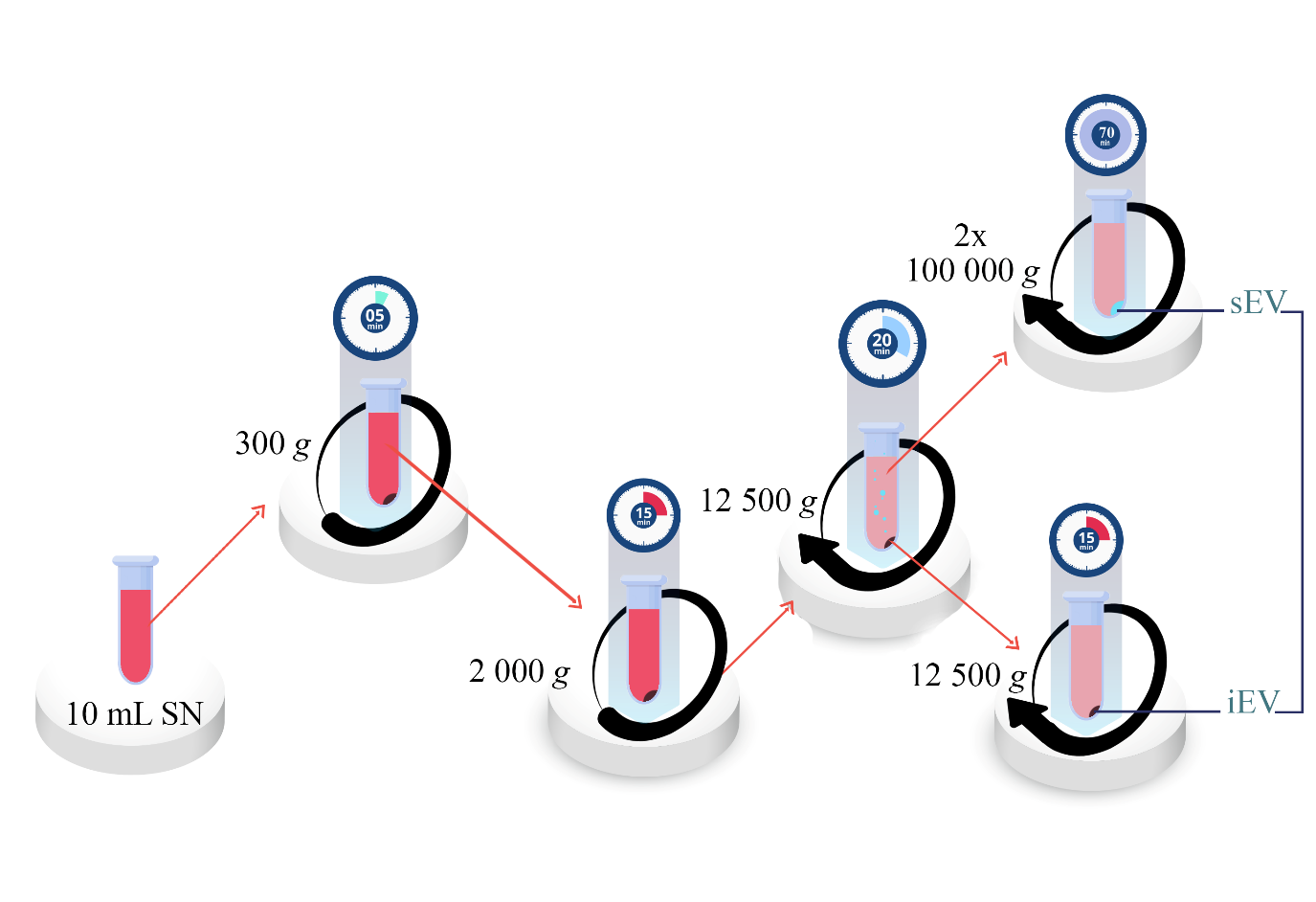

Differential centrifugation was used to isolate extracellular vesicles (EVs) from the medium of serum-starved BeWo cells. The BeWo cell culture supernatant was then centrifuged at 300 g for five minutes at room temperature to remove cell debris. The resulting supernatant was then centrifuged at 2,000 g for 15 minutes to remove apoptotic bodies. The 2,000 g supernatant was then sedimented at 12,500 g for 20 minutes at 16°C using a Z216 MK Microlite centrifuge with a fixed-angle 200.88 rotor (Hermle Labortechnik GmbH, Wehingen, Germany). The sample was subsequently washed with 0.20 µm filtered phosphate-buffered saline (PBS) at 12,500 g for 15 minutes at 16°C. The 12,500 g supernatant was further processed at 100,000 g for 70 minutes at 4 °C (Optima MAX-XP, fixed-angle MLA-55 rotor, Beckman Coulter, Inc., Brea, CA, USA). The sample was washed again with filtered (0.2 µm pore size, Millipore) PBS at 100,000 g for 70 minutes at 4 °C.

**Supplementary Figure 4.**

Characterization of BeWo-derived EVs

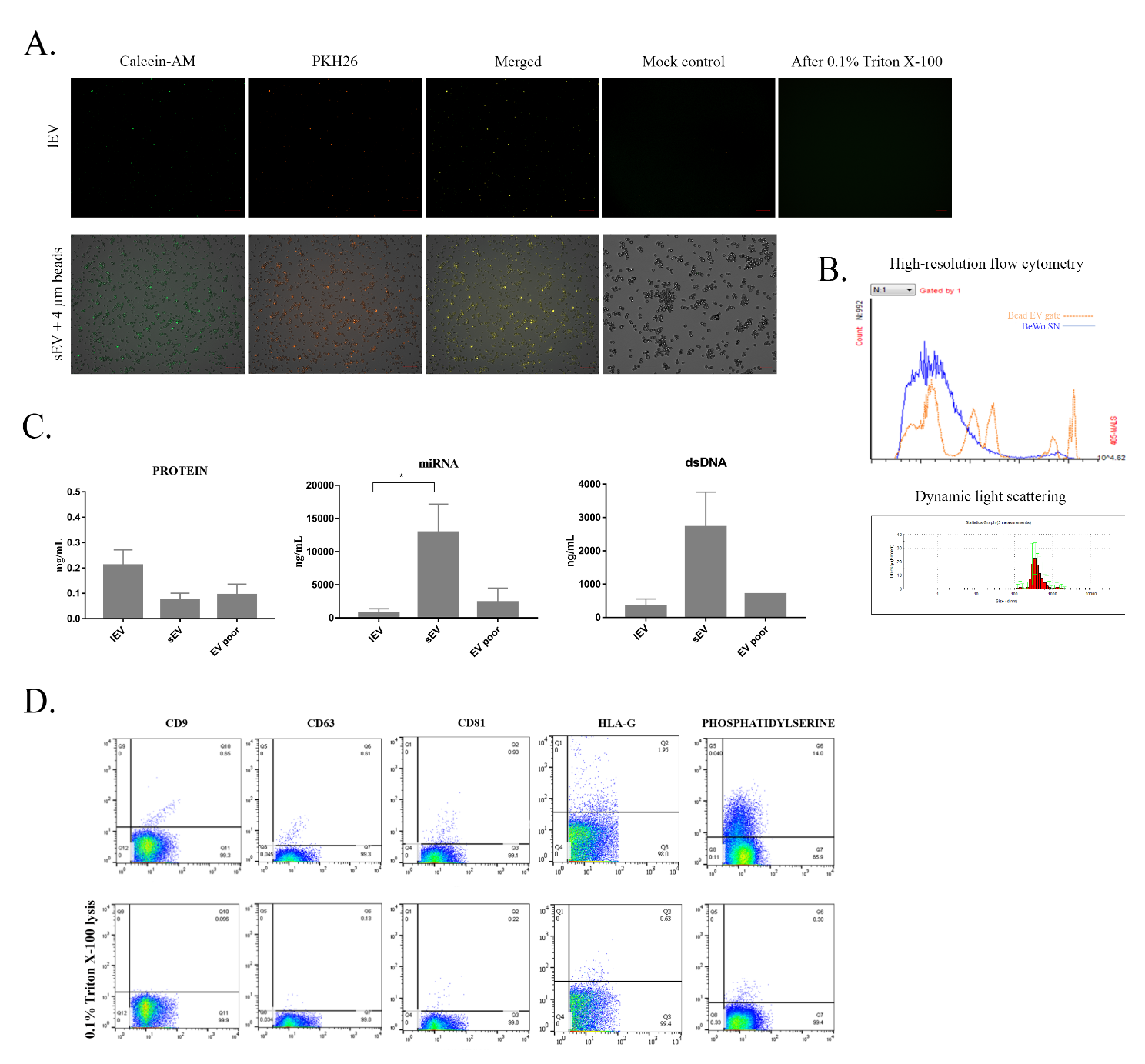

A. High-resolution flow cytometry and dynamic light scattering were used to analyze the size distribution of BeWo-12.5K lEVs, which showed a typical EV size distribution.

B. Representative images of fluorescent microscopy of intermediate-sized and bead-bounded small-sized Calcein-AM-stained BeWo-derived EVs and detergent sensitivity of vesicular structures (scale bar: 1 µm).

C. The bar graphs show the content of proteins, double-stranded DNA (dsDNA), and microRNA (miRNA) in EV preparations. Measurements were performed using a Qubit fluorometer (Qubit Fluorometer, Life Technologies).

D. Representative flow cytometric dot plots of tetraspanins (CD9, CD63, CD81), HLA-G, PS expression on 12.5K lEVs and their corresponding measurements after applying differential detergent lysis with 0.1% Triton X-100.

**Supplementary Figure 5.**

PKH26 staining of 12.5K lEVs

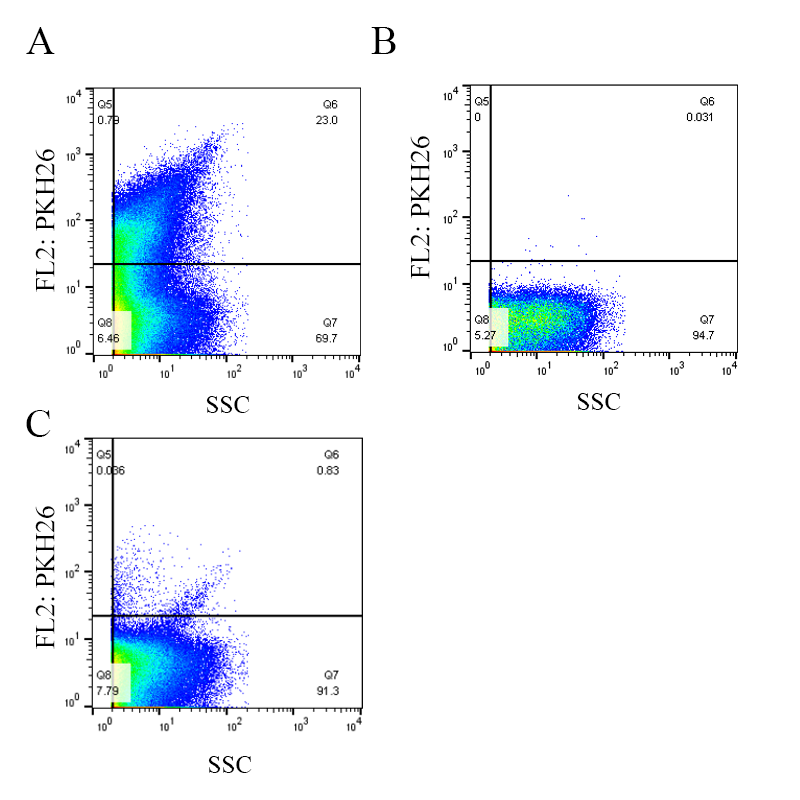

Isolated 12.5K lEVs were labeled with a freshly prepared PKH fluorescent dye solution according to the manufacturer's instructions (Merck, USA). After staining, the labeled vesicles were washed and analyzed by flow cytometry. Staining control (the same staining procedure without adding EV samples) and differential detergent lysis was used for validation.

A) The PKH26 fluorescence of the PKH26-stained 12.5K lEVs was detected within the 12.5K lEV gate.

B) The fluorescence signal of the staining control sample is shown inside the 12.5K lEV gate.

C) PKH26 fluorescence was observed inside the 12.5K lEV gate after lysis with 0.1% Triton X-100 detergent. The dye aggregates and protein complexes did not disappear after lysis.

**Supplementary Figure 6.**

IL-6Rα expression of T lymphocyte subsets

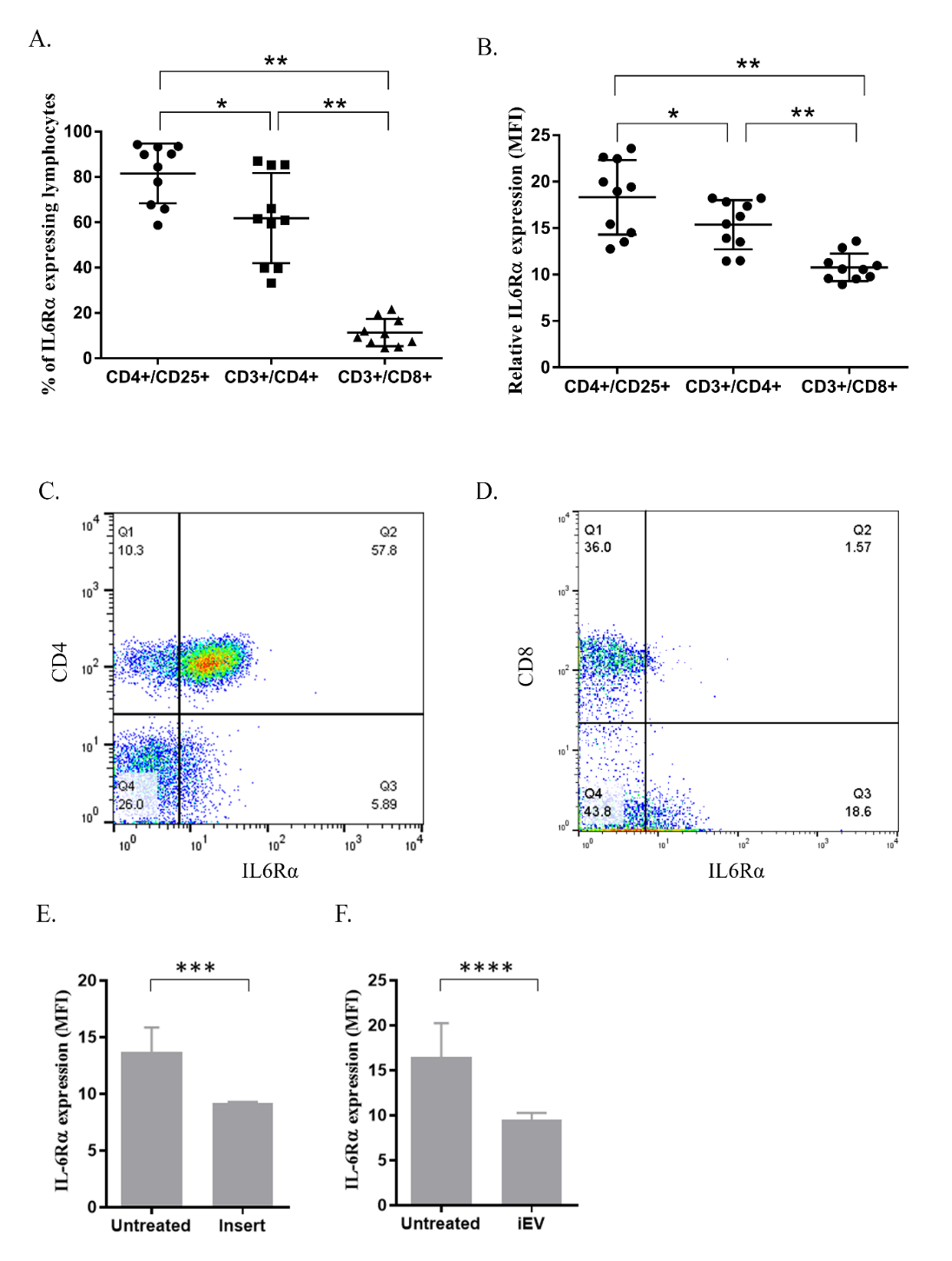

A. Percentage of IL-6Rα expressing T lymphocyte subsets (n=6 pregnant donors)

B. Cell surface expression level of IL-6Rα on lymphocyte subsets (n=6 pregnant donors).

C-D Representative dot plots show the IL-6Rα expression of CD3+/CD4+ and CD3+/CD8+ T cells.

E. IL-6Rα expression of CD4+lymphocytes in BeWo-lymphocyte transwell system.

F. The expression level of IL-6Rα on CD3+CD4+CD25+ cells after treatment with isolated BeWo-derived 12.5K lEVs.

**Supplementary Figure 7**

IL-10 production induced by BeWo-lymphocyte interactions

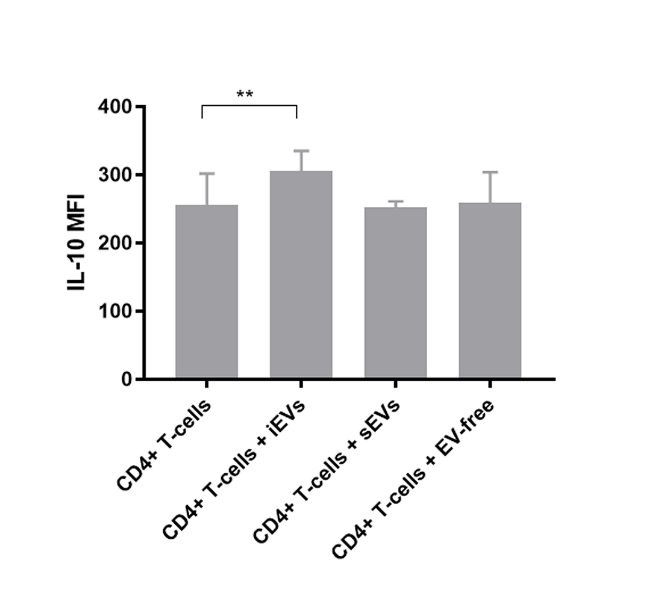

The production of IL-10 by CD4+ T cells was identified by flow cytometry using intracellular staining of IL-10. Small EV fraction (sEV) and EV free cell culture supernatant of BeWo cells (EV-free) were used as biological controls. 12.5K lEVs induced significant IL-10 production in CD4+ T lymphocytes. (** p<0.01, Wilcoxon test). MFI – mean fluorescence intensity (n=3 lymphocyte donors)

**Supplementary table 1** Patient clinical data

|  | Healthy non-pregnant women (n=15) | Healthy pregnant women  (n=37) |
| --- | --- | --- |
| Maternal age (mean ± SD) | 35 ± 6 | 32 ± 4 |
| Gestational age at sampling (mean ± SD) | not applicable | 18 ± 1 |

**Supplementary table 2**

Primers used for qPCR analysis

| GENE | FORWARD PRIMER  OLIGO SEQUENCE (5' TO 3') | REVERSE PRIMER  OLIGO SEQUENCE (5' TO 3') |
| --- | --- | --- |
| **AKT1** | GCACAAACGAGGGGAGTACAT | CCTCACGTTGGTCCACATC |
| **CCL2** | GATCTCAGTGCAGAGGCTCG | TGCTTGTCCAGGTGGTCCAT |
| **CD80** | TTGGATTGTCATCAGCCCTGC | ATTTTCTCCTTTTGCCAGTAG |
| **CSF3R** | CCACGGAGGCAGCTTTAC | AAATCAGCATCCTTTGGGTG |
| **CXCL12** | TCAGCCTGAGCTACAGATGC | CTTTAGCTTCGGGTCAATGC |
| **HPRT** | GGTCAGGCAGTATAATCCAAAG | GTCAAGGGCATATCCTACAAC |
| **IKBKB** | TCCGATGGCACAATCAGGAAAC | TCCAGGCACCACCGCTCTC |
| **IL-1R1** | AGAGGAAAACAAACCCACAAGG | CTGGCCGGTGACATTACAGAT |
| **IL-2RA** | CCAGCTCAGTCCCATCAGAGA | TTCAACGGCGAAATTGCTATT |
| **IL-6** | AGCCACTCACCTCTTCAGAAC | GCCTCTTTGCTGCTTTCACAC |
| **IL-6R** | GACAATGCCACTGTTCACTG | GCTAACTGGCAGGAGAACTT |
| **IL-6ST** | GATGACAAGATGTCCCTCCG | AAA GGACAGGATGTTGCAGG |
| **IL-10** | GTGATGCCCCAAGCTGAGA | CACGGCCTTGCTCTTGTTTT |
| **IL-18R1** | ACGCCGAGTTTGAAGATCAGGGGT | CCCTGGGCAAAATCTCCACAGCA |
| **JAK2** | TTTGGCAACAGACAAATGGA | TGCAGATTTCCCACAAAGTG |
| **JAK3** | GCCTGGAGTGGCATGAGAA | CCCCGGTAAATCTTGGTGAA |
| **LIFR** | TGGAGGGACTGCACAGGTTATTTA | CGGTGACACTGTTAATCGTTTGGT |
| **MAPK1** | CCCAAATGCTGACTCCAAAGC | GCTCGTCACTCGGGTCGTAAT |
| **MAP2K1** | CAATGGCGGTGTGGTGTTC | AGCTCCCTTATGATCTGGTTCC |
| **MAPK3** | ACCTGCGACCTTAAGATTTGT | GAAAAGCTTGGCCCAAGCC |
| **MAPK8** | TGGACTTGGAGGAGAGAACC | ACGATGATGATGGATGCTGA |
| **MAPK14** | AGAAGATGCTTGTATTGGACTCAG | ACAGAAACCAGGTGCTCAGG |
| **MYC** | CACTTTGCACTGGAACTTACAACA | CCCGCGTCGGGAGAGT |
| **MTOR** | CGCTGTCATCCCTTTATCG | ATGCTCAAACACCTCCACC |
| **NFKB1** | TGCCAACAGATGGCCCATAC | TGTTCTTTTCACTAGAGGCACCA |
| **NFKBIA** | CTCCGAGACTTTCGAGGAAATAC | GCCATTGTAGTTGGTAGCCTTCA |
| **OSMR** | ACCTGCCACAGAGTACATGG | GCTCCAAGCTCACAATTCTCCA |
| **PIAS3** | GACTCTCAGCCACTGTTCCC | GCCTCACCAGGTACACAGAC |
| **PPRG** | AAAGAAGCCGACACTAAACC | CTTCCATTACGGAGAGATCC |
| **RAC1** | CCGTGCAAAGTGGTATCCTG | GCTTCTTCTCCTTCAGTTTCTCG |
| **RELA** | TAAGCAGAAGCATTAACTTCTCTGGA | CCTGCTTCTGTCTCTAGGAGAGTA |
| **STAT1** | TGCTCCTTTGGTTGAATCCC | GGAATTTTGAGTCAAGCTGCTG |
| **STAT3** | CCCCATACCTGAAGACCAAGTTTA | CTTCACCATTATTTCCAAACTGCAT |
| **TLR4** | TGGAAGTTGAACGAATGGAATGTG | ACCAGAACTGCTACAACAGATACT |
| **TNFRSF1A** | TGCCTACCCCAGATTGAGAA | ATTTCCCACAAACAATGGAGTAG |
| **TNFRSF1B** | TTCGCTCTTCCAGTTGGACT | CACCAGGGGAAGAATCTGAG |
| **TNFRSF6** | TTATCTGATGTTGACTTGAG | ATTACGAAGCAGTTGAAC |
| **TNFRFS10B** | AAGACCCTTGTGCTCGTTGT | AGGTGGACACAATCCCTCTG |

**Supplementary table 3.**

List of antibodies and dies used in flow cytometry, fluorescent microscopy, western blot and immune electron microscopy

| **Antibody/Dye name** | **Clone*** | **Manufacturer** | **Catalogue number** | **Applied Methods** |
| --- | --- | --- | --- | --- |
| PE anti-human HLA-G | 87G | Biolegend | 335906 | FACS – EV |
| APC anti-human HLA-G | 87G | eBioscience | 17995742 | FACS EV binding |
| FITC Annexin V |  | Biolegend | 640906 | FACS-EV |
| PE Annexin V |  | SONY | 3804540 | FACS-EV |
| PE anti-human CD63 | MEM 259 | Sigma | SAB4700218 | FACS-EV |
| anti-CD63 | H-193 | Sant Cruz Biotech | 15363 | TEM immunogold |
| APC anti-human CD3 | SP  34-2 | BD Biosciences | [557597](http://www.bdbiosciences.com/us/reagents/research/antibodies-buffers/immunology-reagents/anti-non-human-primate-antibodies/cell-surface-antigens/apc-mouse-anti-human-cd3/p/557597) | FACS-lymphocytes |
| PerCP-Cy5.5 anti-human CD4 | RPA-T4 | BD Biosciences | [560650](http://www.bdbiosciences.com/us/applications/research/stem-cell-research/hematopoietic-stem-cell-markers/human/negative-markers/percp-cy55-mouse-anti-human-cd4-rpa-t4/p/560650) | FACS-lymphocytes |
| Pe anti-human CD8 | RPA-T8 | BD Biosciences | [557086](http://www.bdbiosciences.com/us/reagents/research/antibodies-buffers/immunology-reagents/anti-non-human-primate-antibodies/cell-surface-antigens/pe-mouse-anti-human-cd8-rpa-t8/p/557086) | FACS-lymphocytes |
| PerCP-Cy5.5 anti-human CD19 | HIB19 | BD Biosciences | 561295 | FACS-lymphocytes |
| Pe anti-human CD56 | B159 | BD Biosciences | \| [561903](http://www.bdbiosciences.com/us/applications/research/stem-cell-research/hematopoietic-stem-cell-markers/human/negative-markers/pe-mouse-anti-human-cd56-b159/p/561903) | FACS-lymphocytes |
| FITC anti-human CD25 | M-A251 | BD Biosciences | [555431](http://www.bdbiosciences.com/us/applications/research/t-cell-immunology/regulatory-t-cells/surface-markers/human/fitc-mouse-anti-human-cd25-m-a251/p/555431) | FACS-lymphocytes |
| purified anti-human CD95 | DX2 | BD Biosciences | [555670](http://www.bdbiosciences.com/us/applications/research/t-cell-immunology/regulatory-t-cells/surface-markers/human/purified-nale-mouse-anti-human-cd95-dx2/p/555670) | FACS EV binding |
| purified anti-human CD95L | NOK-1 | BD Biosciences | [556372](http://www.bdbiosciences.com/us/applications/research/apoptosis/purified-antibodies/purified-mouse-anti-human-cd178-nok-1/p/556372) | FACS EV binding |
| PKH26 |  | Sigma | P9691 | FACS-EV, Binding Assay, CellDiscoverer7 |
| PKH67 |  | Sigma | PKH67GL | FACS Binding Assay |
| AF488 anti-human CD47 | 1/1A4 | AbD Serotec | MCA2514A488T | FACS – EV |
| Goat anti-Rabbit IgG 10nm gold labelled | polyclonal | Abcam | ab27234 | TEM immunogold |
| Goat anti-mouse 5nm gold labelled | polyclonal | Sigma | G7527 | TEM immunogold |

* In case of antibodies; FACS – EV – flow cytometry of extracellular vesicles; FACS-PBMC –immunophenotyping TEM – transmission electron microscopy

**Supplementary Table 4**

Co-culture system IL-6 pathway analysis

| Gene | Ly | Ly + 12.5K lEV | Ly + BeWo | | Ly insert BeWo |
| --- | --- | --- | --- | --- | --- |
|  | Relative expression | | | | |
| *STAT1* | 265.8 ± 128 | 3.82 ± 0.38 | | 308.4 ± 148 | 990 ± 75.2 |
| *PIAS3* | 44.7 ± 0.07 | 4.48 ± 0.01 | | 1.10 ± 0.37 | Not detectable |
| *AKT1* | 16.76 ± 0.68 | 23.30 ± 0.42 | | 17.39 ± 3.15 | 21.50 ± 4.02 |
| *RELA* | 0.36 ± 0.01 | 4.04 ± 2.06 | | 0.16 ± 0.09 | 0.42 ± 0.07 |
| *NFKBIA* | 67.80 ± 48.6 | 268.2 ± 2.6 | | 26.61 ± 15.4 | 111.1 ± 47.5 |
| *CXCL12* | 2.03 ± 0.18 | 1.62 ± 0.24 | | 1.35 ± 1.14 | 2.02 ± 0.26 |
| *IL1R1* | 9.61 ± 0.53 | 7.51 ± 0.88 | | 6.63 ± 0.89 | 12.10 ± 0.75 |
| *IL6ST* | 399.5 ± 64.5 | 374.2 ± 28.21 | | 159.7 ± 37.1 | 361.1 ± 156 |
| *OSMR* | 11.12 ± 0.61 | 10.51 ± 0.22 | | 7.88 ± 1.71 | 12.06 ± 0.31 |
| *TNFRSF10B* | 2.05 ± 0.01 | 2.07 ± 0.14 | | 1.53 ± 0.23 | 2.28 ± 0.10 |
| *STAT3* | 627.4 | 432.8 ± 84.3 | | 239.1 ± 130 | Not detectable |
| *IL10* | 2.04 ± 0.18 | 1.98 ± 0.09 | | 1.49 ± 0.3 | 2.18 ± 0.7 |
| *IL2RA* | 64.31 ± 7.5 | 55.5 ± 5.0 | | 64.14 ± 2.1 | 79.11 ± 23.3 |
| *CD80* | 1.18 ± 0.01 | 1.10 ± 0.48 | | 0.94 ± 0.26 | 1.63 ± 0.10 |
| *JAK2* | 0.96 ± 0.02 | 0.85 ± 0.13 | | 0.66 ± 0.12 | 1.11 ± 0.04 |
| *JAK3* | 0.23 ± 0.01 | 0.28 ± 0.01 | | 0.22 ± 0.05 | 0.40 ± 0.01 |
| *MAPK1* | 2.03 ± 0.13 | 2.14 ± 0.55 | | 3.15 ± 0.54 | 5.68 ± 1.03 |
| *MAPK14* | 3.67 ± 0.24 | 4.39 ± 0.35 | | 2.38 ± 0.98 | 5.46 ± 0.04 |
| *RAC1* | 93.1 ± 43.3 | 209.1 ± 24.4 | | 134.0 ± 20 | 312.9 ± 32.9 |

**Supplementary methods:**

Dynamic light scattering measurement of EVs

The size distribution of EVs were characterized by dynamic light scattering on a Zetasizer Nano S instrument (Malvern Instruments Ltd, Malvern, UK). From the intensity fluctuations of a 633-nm laser light scattered at high angle from the freely moving suspended particles their diffusion constant was obtained. Size distribution was calculated by using the Stokes-Einstein equation by the built-in algorithms of the instrument’s software. Z-average values are displayed, which represent the primary and most stable parameter produced by dynamic light scattering technique and recommended for quality control reports (ISO 22412:2008). Z-average value represents a good approximation of hydrodynamic diameter of well dispersed particles. Polydispersity index (PDI) is an estimate of the width of the distribution which is calculated from the cumulants analysis also.

Light scattering was measured at 20 ± 1°C. Vesicle (refractive index: 1.380) was suspended in 0.2 µm filtered PBS. The viscosity of PBS at 20 °C was 1.0041 cP and the refractive index was 1.330. The sample was equilibrated at 20 °C for 1 min.
